# Light Pollution Increases West Nile Virus Competence in a Ubiquitous Passerine Reservoir Species

**DOI:** 10.1101/269209

**Authors:** M. E. Kernbach, J. M. Miller, R. J. Hall, T. R. Unnasch, N. D. Burkett-Cadena, L. B. Martin

## Abstract

**One sentence summary:** Light pollution increases host infectiousness.

**Abstract:** Light pollution is a growing problem, but its impacts on infectious disease risk have not been considered. Previous research has revealed that dim light at night (dLAN) dysregulates various immune functions and biorhythms, which hints that dLAN could change the risk of disease epidemics. Here, we demonstrate that dLAN enhances infectiousness of the house sparrow (*Passer domesticus*), an urban-dwelling avian host of West Nile virus (WNV). Sparrows exposed to dLAN maintained viral titers above the transmission threshold to a biting vector (10^5^ plaque-forming units) for two days longer than controls but did not die at higher rates. A mathematical model revealed that such effects could increase WNV outbreak potential by ~41%. dLAN likely affects other host and vector traits relevant to transmission, so additional research is needed to advise management of zoonotic diseases in light polluted areas.

## Main Text

Among the many anthropogenic changes that impact humans and wildlife, one of the most pervasive but least understood is light pollution (*1*). Light pollution takes many forms (i.e. sky glow, light clutter, glare; 2), but dim light at night (dLAN) is exceptionally common and important in both urban centers and non-urban areas including farms, airports, warehouses, and wherever lighting is necessary for human activities at night (*Fig. 1*). Indeed, greenspaces near roadways experience extensive dLAN exposure from billboards, street lamps, residential lighting, and headlights emanating from passing vehicles (*3*).

**Fig. 1.**
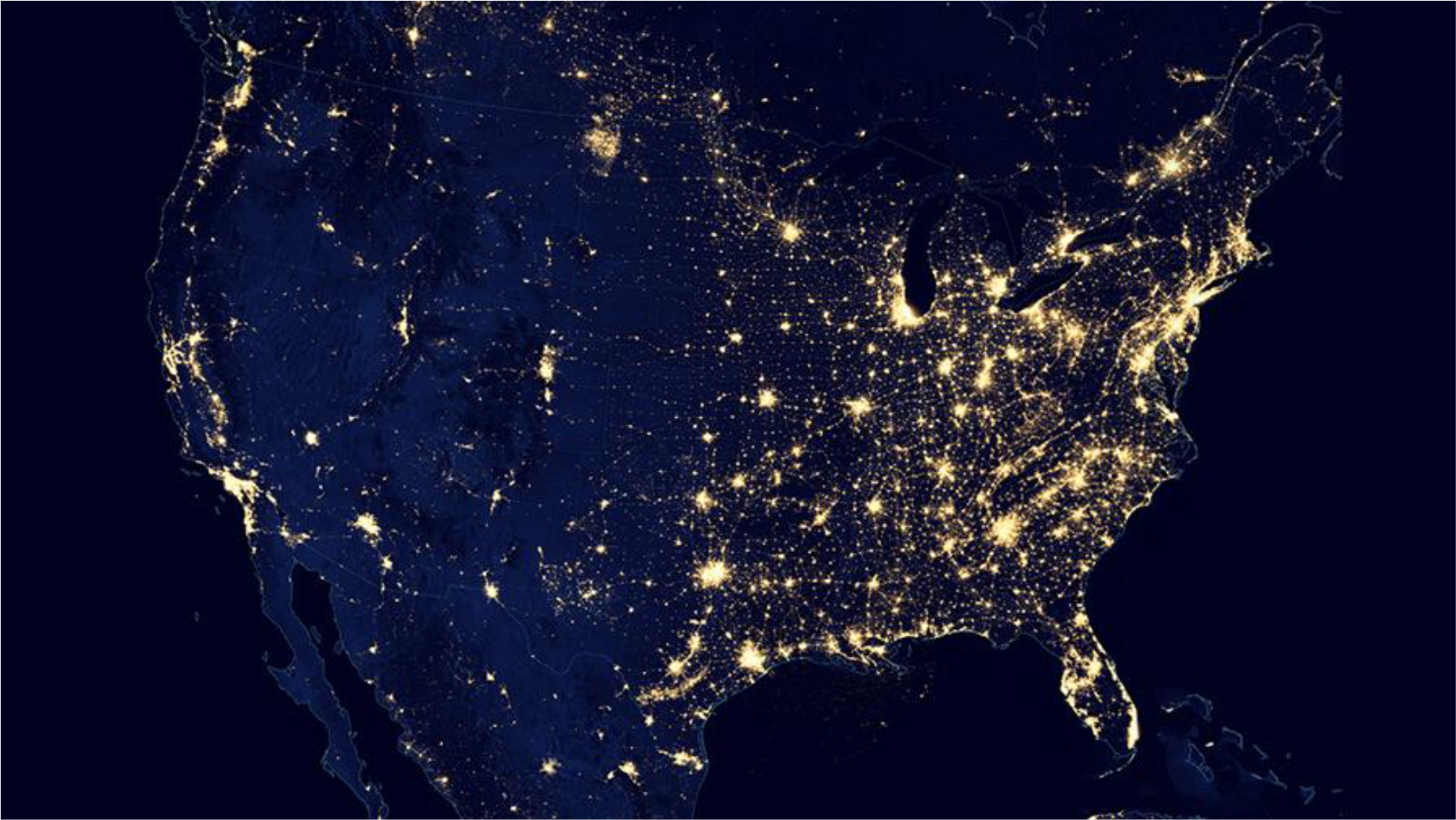
Suomi NPP satellite derived VIIRS image of artificial nighttime illumination across the United States in 2012. Photo credit to NASA Earth Observatory. *Light Pollution*. Digital Image. *NASA Earth Observatory*. n.p, n.d. Web 15 Jan 2018 <https://earthobservatory.nasa.gov/IOTD/view.php?id=84722>

Although dLAN is widespread, its effects on human and wildlife health have been under-studied. Early research on human health found that individuals working throughout the night routinely suffer higher rates of Type II diabetes, heart conditions and other non-infectious maladies compared to day-working staff (*4*). The recent switch to energy efficient night-lighting, which has converted incandescent bulbs to short wavelength cool white LEDs, may increase health risk further. In domesticated rodents, such short wavelength light has been linked to metabolic dysregulation, immunosuppression, and the development of some cancers (*4*). Levels of blue light as low as 0.2 lux can suppress melatonin secretion in humans (*5, 6*), and in wildlife, comparable forms of dLAN alter many behavioral, life history, and physiological traits (*7, 8*).

Despite the diverse and strong effects of dLAN, no study has yet investigated whether and to what degree it might affect infectious disease risk. This is surprising given the pervasiveness of light pollution, especially because hosts and vectors have evolved to use light cues to coordinate multiple daily and seasonal rhythms (*9, 10*). Light is among the most reliable environmental cues, and light regimes induce variation in the timing of animal activities as well as temporal fluctuations in immune defenses and other factors that influence risk of infection (*11*). Our goal was to discern whether light pollution, and dLAN specifically, could alter zoonotic disease risk for humans and wildlife by changing the ability of a reservoir host to amplify virus for subsequent uptake by a vector. Heterogeneity in such an ability, which we term host competence (*12–14*), is mediated by physiological pathways sensitive to light signals (*11*) and those coordinating organismal responses to stressors, namely the glucocorticoids (*15*). Melatonin and glucocorticoids have profound effects on both host behaviors that affect parasite exposure risk and anti-vector defenses (*16–17*).

In the present study, we investigated dLAN effects on WNV infections in house sparrows (*Passer domesticus*) because they are among the most common birds in light-polluted areas (*18*). They are also among the more competent species for WNV (*19*), and a close commensal of humans. We chose WNV for two reasons: i) >46,000 cases of human disease have been reported across the US since its introduction to New York in 1999 (*20*), and ii) it has decimated avian populations, particularly corvids and other passerine species that commonly occupy light-polluted habitats (*21*). In previous work with WNV, we found that corticosterone, the aforementioned avian stress hormone, enhances WNV competence by elevating attractiveness to *Culex* mosquito vectors for blood-feeding (*22*) and increasing WNV viremia above the transmission-to-vector threshold (10^5^ log plaque-forming units (PFUs) (supp. material)). We thus designed our study to determine whether any effects of dLAN on WNV competence were mediated by corticosterone dysregulation.

To assess the effects of dLAN on house sparrow competence for WNV, we exposed wild-caught birds to dLAN (12h light: 12h 8 lux dim light) or natural light conditions (12h light: 12h dark) for 1-3 weeks before and after exposure to 10^1^ (PFUs) of WNV, NY 1999 strain (*13*; dLAN exposure duration was not impactful in models, so it is not further addressed below). Following WNV exposure, we sampled serum on days 2, 4, 6, and 10 to quantify WNV viremia in circulation (*13*), and we measured body mass (to 0.1g prior to and on each blood sampling day) to assess effects on individual health. We detected a significant effect of dLAN on WNV viremia in house sparrows (dLAN x time: F_4,124_ = 2.9, P = 0.023; Figure 2A). Over the course of the first four days post-exposure (dpe; all animals in both groups became infected), both dLAN and control birds had comparable viral titers, but 6 days post-exposure (dpe), dLAN individuals had significantly higher viral titers than control individuals (β = 1.7 +/−0.60, t = −2.7, P = 0.006). Previous studies have found that the minimum circulating viral titer needed to transmit WNV to vectors is ~10^5^ PFUs (horizontal dashed line in Fig. 2A).

**Fig. 2.**
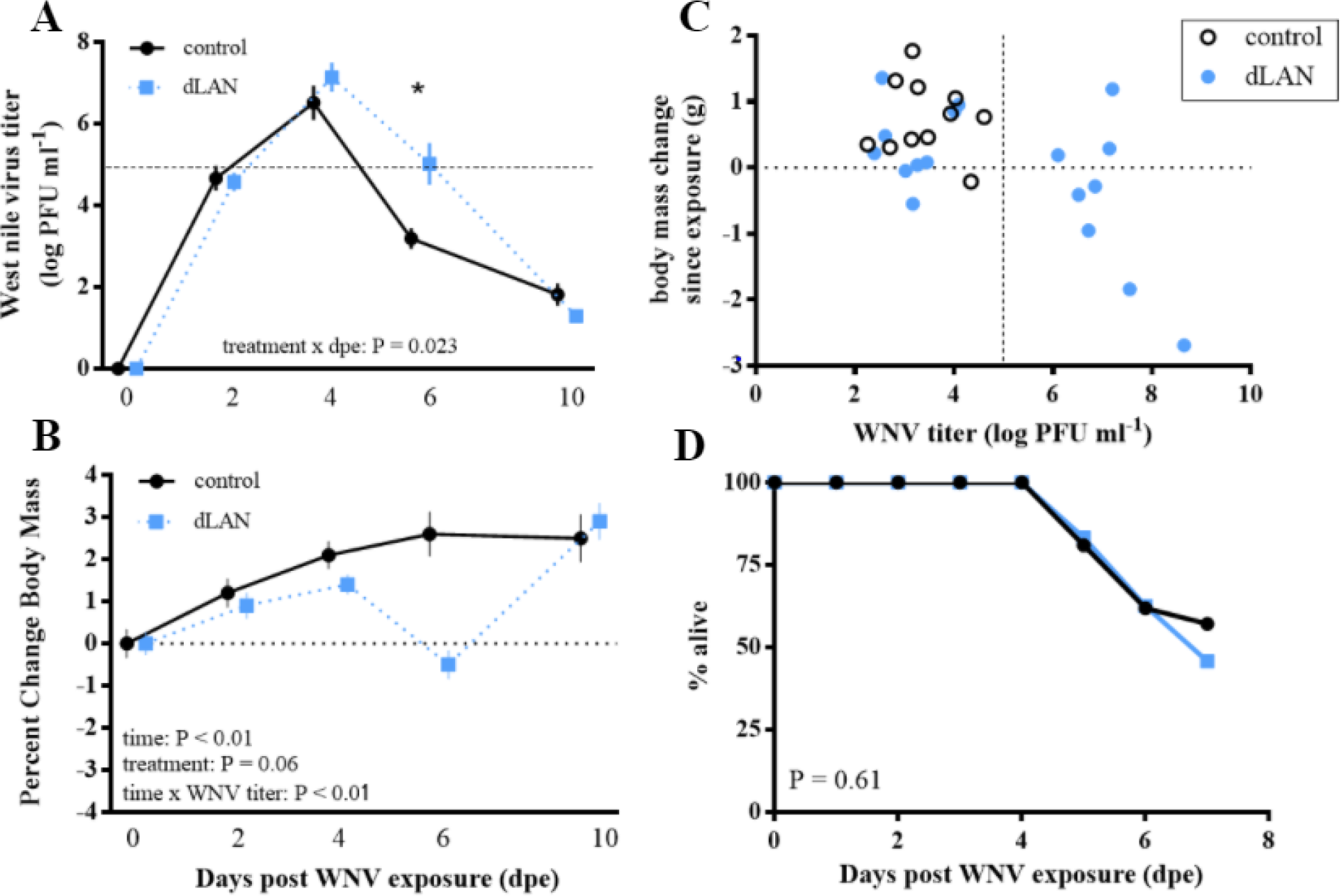
Effects of experimental West Nile virus exposure and impacts on house sparrows (*Passer domesticus*) exposed to dim light at night (dLAN; 8 lux during night hours for 2-3 weeks prior to WNV exposure) versus controls (animals kept on 12h light: 12 dark for duration of experiment). Blue points and lines signify dLAN-exposed individuals, and black points and solid lines signify controls. (A) Individuals exposed to dLAN had significantly higher viral titers on d6 post exposure, indicated by the asterisk. The horizontal dashed light represents the transmission threshold, or the minimum amount of virus in circulation required to transmit WNV to a vector (i.e. 10^5^ PFUs). (B) Effects of WNV and dLAN on change in body mass throughout the course of WNV infection. On d6, dLAN exposed individuals loss appreciable mass whereas controls continued to gain body mass. C) Relationship between WNV titer and body mass change on d6 post WNV exposure. The vertical dashed line represents the WNV transmission threshold; individuals to the right of this dashed line are infectious to mosquitoes, and individuals to the left of this dashed line are not. Only dLAN exposed individuals were infectious on d6. D) No effect of dLAN on WNV-associated mortality. Although some dLAN exposed birds lost more mass and maintained higher viral titers for longer periods of time than controls (in (B)), mortality rate did not differ between groups.

We then asked whether dLAN might modify the capacity of individuals to transmit WNV to biting vectors. When we queried whether dLAN generally affected WNV tolerance (in the form of defense of body mass sensu *23*; % change relative to pre-WNV exposure values), we found no effect across the entire post-exposure period (dLAN x integrated WNV titer: F_1,34_ = 1.3, P = 0.26). However, birds with the highest WNV titers overall lost more body mass than birds with lower cumulative titers (integrated WNV titer: F_1,34_ = 6.6, P = 0.02). Subsequently, we assessed directly whether body mass changed differently post-infection in dLAN and control birds. We found that birds in the control group tended to gain mass post-infection, whereas body mass reached a nadir in dLAN birds on d6 post-infection (Fig. 2B). When we analyzed how body mass and WNV titer were related over the infectious period, we found that the interaction varied with days post-exposure (dpe x dLAN x WNV titer: F_3,83_ = 3.11, P = 0.03). This three-way interaction was driven by WNV x dLAN effects on d6 (β = 1.3 +/−0.60, t = 2.1, P = 0.04; Fig. 2C): at this time, only dLAN-exposed birds (~1/2 of dLAN birds) maintained WNV titers above the transmission threshold; no control birds were infectious on d6. Given the stronger effects of WNV on body mass loss in the dLAN versus control group, one might expect greater mortality in the former. However, we found no effect of dLAN on survival of WNV infection postexposure (χ^2^_1_ = 0.26, P = 0.61; Fig. 2D). About 60% of birds in each group survived to d6.

The second goal of our study, to evaluate one putative mechanism whereby dLAN affected WNV viremia, revealed that dLAN modestly affected CORT levels but this variation did not predict any aspect of WNV competence (see supplemental material). To evaluate the epidemiological implications of dLAN on WNV risk in a wild bird population (Fig. 3), we calculated the relative change in the pathogen basic reproductive number, R_0_, in the presence and absence of dLAN effects based on a single host, single vector model of WNV transmission using parameters derived from literature and these experiments (*24*). We found that dLAN-induced changes in the avian host’s infectious period increased R_0_ by 41% (Supplementary Material). Assuming that dLAN had no other effects on avian or vector competence and abundance, we estimated that dLAN would increase R_0_ from 8.95 to 12.66 in a WNV-naive songbird community where House Sparrows are the dominant species (Supplementary Material). Our results motivate further investigation of the ecological and molecular mechanisms whereby dLAN alters zoonotic risk. In addition to effects observed here, light pollution might alter vector and host diversity as well as the nature and timing of their interactions (over days and seasons; *25*). Most WNV vectors, for instance, take blood meals at dusk and dawn (*26*); with dLAN, the blood-meal feeding window might be extended, or vectors might arouse too early to find a blood meal before they are forced to forage on nectars or other substrates just to remain viable. The pineal-derived hormone, melatonin, which coordinates circadian oscillators that reside in cells throughout the body, often regulates such behaviors (*27*). Dysregulation of melatonin synthesis and secretion, which has been observed in response to dLAN in the lab (*4*), could have complex effects on WNV dynamics, particularly as vectors too rely on melatonin for temporal coordination of behaviors. In urban areas, comparatively warmer and more stable temperatures and numerous habitats for breeding could augment mosquito populations. Conversely, urban vectors might lay few eggs collectively because of difficulties in synchronizing blood-meal searching with periods of optimal host availability. Coupled with the ability of melatonin to modify host and vector immune functions (*28*), it is very difficult to predict now how dLAN will truly affect disease risk in natural settings.

As we start to investigate dLAN effects on infectious disease risk, zoonotic and otherwise, it will be important to study how the lighting spectra we currently favor (and when and where we use lighting) can be adjusted to mitigate risk. Motion-activated or directionally-targeted light sources can be substituted for the constant illumination practices, and lighting overall could also be reduced when it would have the greatest impacts on wildlife (i.e., migrations, breeding seasons). A good example to emulate comes from the International Dark-Sky Association, which has led efforts to eliminate lighting in tall urban buildings during avian migrations; this practice reduces the extensive window strikes that occur during those periods (*2*). An equivalent example for vectored-disease in the southeastern US would entail a diminution in lighting of vulnerable areas during the height of arbovirus transmission season (e.g., late Fall; *29*). Additional mitigation opportunities likely reside in the advent of new technologies detectable by human, but less so wildlife, vision (e.g., high-wavelength (red) wavelengths versus the broad-spectrum options typically used; *30*).

## Acknowledgments

We thank Erik Hofmeister for sharing the West Nile virus NY’99 strain, and we thank members of the Martin lab for help on experimental methods and protocols.

## Author contributions

M. E. Kernbach contributed to conceptualization, data curation, methodology, investigation, project administration, and writing-original draft; J. M. Miller contributed to data curation, project administration, and investigation; R. J. Hall contributed to conceptualization, formal analysis, visualization, writing-original draft; T. R. Unnasch contributed to conceptualization, funding acquisition, methodology, resources, and supervision; N. D. Burkett-Cadena contributed to formal analysis, investigation, and methodology; L. B. Martin contributed to conceptualization, data curation, formal analysis, funding acquisition, investigation, methodology, project administration, resources, supervision, and writing-original draft.

## Data and materials availability

All data code and material used to produce this manuscript will be uploaded to Dryad upon acceptance.

## References and Notes

1. M. Leu, S. Hanser, S. Knick, The Human Footprint in the West: A Large-Scale Analysis of Anthropogenic Impacts. Ecological Applications, 18(5), 1119–1139 (2008).

2. “International Dark-Sky Association’s Practical Guide 1: Introduction to Light Pollution”. IDA. (2009) [no author]

3. P. Cinzano, F. Falchi, C. D. Elvidge, The first World Atlas of the artificial night sky brightness. Monthly Notices of the Royal Astronomical Society, 328(3), 689–707 (2001)

4. K. J. Navara, R. J. Nelson, The dark side of light at night: physiological, epidemiological, and ecological consequences. Journal of Pineal Research, 43(3), 215–224 (2007).

5. K. Thapan, J. Arendt, D. J. Skene, An action spectrum for melatonin suppression: evidence for a novel non-rod, non-cone photoreceptor system in humans. The Journal of Physiology, 535(1), 261–267 (2001).

6. S. M. Pauley, Lighting for the human circadian clock: recent research indicates that lighting has become a public health issue. Medical Hypotheses, 63(4), 588–596 (2004).

7. D. M. Dominoni, The effects of light pollution on biological rhythms of birds: an integrated, mechanistic perspective. Journal of Ornithology, 156(1), 409–418 (2015).

8. B. E. Witherington, E. R. Martin, “Understanding, assessing, and resolving light-pollution problems on sea turtle nesting beaches” (FMRI Technical Report TR-2, 2003).

9. M. H. Hastings, J. Herbert, N. D. Martensz, A. C. Roberts, Annual reproductive rhythms in mammals: mechanisms of light synchronization. Annals of the New York Academy of Sciences, 453(1), 182 (1985)

10. U. Schibler, The daily rhythms of genes, cells and organs. EMBO Reports, 6(S1), S9–-S13 (2005).

11. T. A. Bedrosian, L. K. Fonken, J. C. Walton, R. J. Nelson, Chronic exposure to dim light at night suppresses immune responses in Siberian hamsters. Biology Letters, 7(3):468–71 (2011).

12. D. Barron, S. Gervasi, J. Pruitt, L. B. Martin, Behavioral competence: how host behaviors can interact to influence parasite transmission risk. Current Opinion in Behavioral Sciences, 6, 35–40 (2015).

13. S. S. Gervasi, S. C. Burgan, E. Hofmeister, T. R. Unnasch, L. B. Martin, Stress hormones predict a host superspreader phenotype in the West Nile virus system. Proceedings of the Royal Society of London B: Biological Sciences, 284(1859) (2017).

14. S. H. Paull, S. Song, K. M. McClure, L. C. Sackett, A. M. Kilpatrick, P. T. Johnson, From superspreaders to disease hotspots: linking transmission across hosts and space. Frontiers in Ecology and the Environment, 10(2), 75–82 (2012).

15. L. B. Martin, S. C. Burgan, J. S. Adelman, S. S. Gervasi, Host Competence: An Organismal Trait to Integrate Immunology and Epidemiology. Integrative and Comparative Biology, icw064 (2016).

16. J. Q. Ouyang, M. de Jong, M. Hau, M. E. Visser, R. H. A. van Grunsven, K. Spoelstra, Stressful colours: corticosterone concentrations in a free-living songbird vary with the spectral composition of experimental illumination. Biology Letters, 11(8) (2015).

17. R. M. Sapolsky, L. M. Romero, A. U. Munck, How Do Glucocorticoids Influence Stress Responses? Integrating Permissive, Suppressive, Stimulatory, and Preparative Actions*. Endocrine Reviews, 21(1), 55–89 (2000)

18. D. E. Chamberlain, M. P. Toms, R. Cleary-McHarg, A. N. Banks, House sparrow (Passer domesticus) habitat use in urbanized landscapes. Journal of Ornithology, 148(4), 453–462 (2007)

19. N. Komar, S. Langevin, S. Hinten, N. Nemeth, E. Edwards, D. Hettler, B. Davis, R. Bowen, M. Bunning, Experimental infection of North American birds with the New York 1999 strain of West Nile virus. Emerging Infectious Diseases, 9(3), 311–322 (2003).

20. “West Nile virus disease cases and deaths reported to CDC by year and clinical presentation, 1999-2016” ArboNET, Arboviral Diseases Branch, Centers for Disease Control and Prevention (2016). [no author]

21. P. P. Marra, S. Griffing, C. Caffrey, M. A. Kilpatrick, R. McLean, C. Brand, E. Saito, A. P. Dupuis, L. Kramer, R. Novak, West Nile Virus and Wildlife. BioScience, 54(5), 393–402 (2004).

22. S. Gervasi, S. Burgan, N. Burkett-Cadena, A. Schrey, H. Hassan, T. Unnasch, L. Martin, Vector preference and host defenses in the West Nile virus system: A role for avian stress hormones? Integrative and Comparative Biology, 56, E74 (2016).

23. L. Råberg, D. Sim, A. F. Read, Disentangling Genetic Variation for Resistance and Tolerance to Infectious Diseases in Animals. Science, 318(5851), 812 LP–814 (2007).

24. M. J. Wonham, T. De-Camino-Beck, M. A. Lewis, An epidemiological model for West Nile virus: invasion analysis and control applications. Proceedings of the Royal Society of London B: Biological Sciences, 271(1538), 501–507 (2004).

25. V. O. Ezenwa, M. S. Godsey, R. J. King, S. C. Guptill, Avian diversity and West Nile virus: testing associations between biodiversity and infectious disease risk. Proceedings. Biological Sciences, 273(1582), 109–17 (2006).

26. H. M. Savage, M. Anderson, E. Gordon, L. Mcmillen, L. Colton, M. Delorey, … M. Godsey, Host-seeking heights, host-seeking activity patterns, and West Nile virus infection rates for members of the Culex pipiens complex at different habitat types within the hybrid zone, Shelby County, TN, 2002 (Diptera: Culicidae). Journal of Medical Entomology, 45(2), 276–288 (2008).

27. E. Gwinner, I. Benzinger, Synchronization of a circadian rhythm in pinealectomized European starlings by daily injections of melatonin. Journal of Comparative Physiology, 127(3), 209–213 (1978).

28. A. Carrillo-Vico, P. J. Lardone, N. Álvarez-Sánchez, A. Rodríguez-Rodríguez, J. M. Guerrero, Melatonin: Buffering the Immune System. International Journal of Molecular Sciences, 14(4), 8638–8683 (2013).

29. C. G. M. Blackmore, L. M. Stark, W. C. Jeter, R. L. Oliveri, R. G. Brooks, L. A. Conti, S. T. Wiersma, Surveillance results from the first West Nile transmission season in Florida, 2001. The American Journal of Tropical Medicine and Hygiene, 69(2) (2003).

30. A. Kelber, M. Vorobyev, D. Osorio, Animal colour vision - behavioural tests and physiological concepts. Biological Reviews, 78(1), 81–118 (2003).

31. J. Adelman, L. Kirkpatrick, J. Grodio, D. Hawley, House Finch Populations Differ in Early Inflammatory Signaling and Pathogen Tolerance at the Peak of Mycoplasma gallisepticum Infection. The American Naturalist, 181(5), 674–689 (2013).

32. P. T. J. Johnson, J. R. Rohr, J. T. Hoverman, E. Kellermanns, J. Bowerman, K. B. Lunde, Living fast and dying of infection: host life history drives interspecific variation in infection and disease risk. Ecology Letters, 15(3), 235–242 (2012).

33. A. L. Liebl, T. Shimizu, L. B. Martin, Covariation among glucocorticoid regulatory elements varies seasonally in house sparrows. General and Comparative Endocrinology, 183, 32–37 (2013).

34. L. B. Martin, L. Kidd, A. L. Liebl, C. A. Coon, Captivity induces hyper-inflammation in the house sparrow (Passer domesticus). Journal of Experimental Biology, 214(15), 2579–2585 (2011).

35. M. J. Wonham, T. De-Camino-Beck, M. A. Lewis, An epidemiological model for West Nile virus: invasion analysis and control applications. Proceedings of the Royal Society of London B: Biological Sciences, 271(1538), 501–507 (2004).

36. E. C. Uttah, G. N. Woekm, C. Okonofua, The abundance and biting patterns of Culex quinquefasciatus Say (Culicidae) in the coastal region of Nigeria. ISRN Zoology. 2013(640691), (2013).

37. M. J. Turell, M. L. O’Guinn, D. J. Dohm, J. W. Jones, Vector competence of North American mosquitoes (Diptera: Culicidae) for West Nile virus. Journal of Medical Entomology, 38(2), 130–134 (2001).

38. M. R. David, G. S. Ribeiro, R. M. de Freitas, Bionomics of Culex quinquefasciatus within urban areas of Rio de Janeiro, Southeastern Brazil. Journal of Public Health, 46(5), 858–65 (2012).

39. S. L. Richards, C. N. Mores, C. C. Lord, W. J. Tabachnick, Impact of extrinsic incubation temperature and virus exposure on vector competence of Culex pipiens quinquefasciatus Say (Diptera: Culicidae) for West Nile virus. Vector Borne and Zoonotic Diseases (Larchmont, N.Y.), 7(4), 629–636 (2007).

40. M. F. Sallam, S. R. Michaels, C. Riegel, R. M. Pereira, W. Zipperer, B. G. Lockaby, P. G. Koehler, Spatio-temporal distribution of vector-host contact (VHC) ratios and ecological niche modeling of the West Nile virus mosquito vector, Culex quinquefasciatus, in the city of New Orleans, LA, USA. International Journal of Environmental Research and Public Health, 14(8), 892 (2017).

41. M. Turell, M. O’Guinn, J. Oliver, Potential for New York mosquitoes to transmit West Nile virus. Am. J. Trop. Med. Hyg. 62, 413–414 (2000).

